# Saltational episodes of reticulate evolution in the *Drosophila saltans* species group

**DOI:** 10.1101/2023.10.09.561511

**Authors:** Carolina Prediger, Erina A. Ferreira, Samara Videira Zorzato, Aurélie Hua-Van, Lisa Klasson, Wolfgang J. Miller, Amir Yassin, Lilian Madi-Ravazzi

## Abstract

Phylogenomics reveals reticulate evolution to be widespread across taxa, but whether reticulation is due to low statistical power or it is a true evolutionary pattern remains a field of investigation. Here, we investigate the phylogeny and quantify reticulation in the *Drosophila saltans* species group, a Neotropical clade of the subgenus *Sophophora* comprising 23 species whose relationships have long been problematic. Phylogenetic analyses revealed conflicting topologies between the X chromosome, autosomes and the mitochondria. We extended the ABBA-BABA test of asymmetry in phylogenetic discordance to cases where no “true” species tree could be inferred, and applied our new test (called 2A2B) to whole genome data and to individual loci. We used four strategies, two of which consisted of windows from pseudo-reference genomes aligned to either an outgroup or ingroup species, and two based on our new assemblies using either conserved genes or ≥50 kb-long syntenic blocks with conserved collinearity across Neotropical *Sophophora*. Evidence for reticulation varied among the strategies, being lowest in the synteny-based approach, where it did not exceed ∼7% of the blocks in the most conflicting species quartets. High incidences of low phylogenetic resolution (polytomy) were restricted to three nodes on the tree, that coincided with major paleogeographical events in South America. Our results identify possible technical biases in quantifying reticulate evolution and indicate that episodic rapid radiations have played a major role in the evolution of a largely understudied Neotropical clade.

## Introduction

Knowledge of phylogenetic relationships among species is a requirement for many evolutionary studies. However, it is often difficult to reconstruct well-resolved bifurcating trees for some clades. This could either be due to the lack of signal in the evaluated data, a condition known as “soft polytomy”, or due to persistent phylogenetic conflicts among datasets leading to “hard polytomies” and reticulate patterns of interspecific relationships. A plethora of biological processes could cause such conflicts, including incomplete lineage sorting (Maddison 1989; Maddison 1997; Walsh et al. 1999; Townsend et al. 2012), horizontal gene transfer, introgression and hybridization (Schrempf and Szöllősi 2020), and adaptive radiations (Glor 2010). Phylogenetic conflict may also be caused by technical errors, such as, sequencing error, contamination, wrong model selection and general lack of quality control (Philippe et al. 2011). Recent advances in genomic analyses have significantly reduced such errors and, in a wide range of taxa, increased the number of analyzed genes hence helping to resolve early conflicting topologies. However, in many other cases, whole genome analyses demonstrated persistent phylogenetic conflicts (e.g., in plants (Wickett et al. 2014; Gagnon et al. 2022), birds (Suh 2016), sponges and ctenophores (Philippe et al. 2009; Pick et al. 2010; Chang et al. 2015; Whelan et al. 2015; Simion et al. 2017), mammals (Morgan et al. 2013; Romiguier et al. 2013; Doronina et al. 2015), amphibians (Hime et al. 2021), and insects (Owen and Miller 2022)).

Of the different processes that can reduce phylogenetic resolution, introgression and hybridization have attracted much attention, and led to the development of a number of bioinformatic tools and tests that quantify their extent across the genome (Durand et al. 2011; Pease and Hahn 2015; Malinsky et al. 2021). Site-based methods usually count the number of bi-allelic sites supporting each of three possible topologies in a species triplet with an outgroup (Figure 1A). Comparisons between the proportions of the three topologies can yield one of four possible outcomes (Figure 1B): (i) complete trifurcation, all topologies are equally encountered; (ii) incomplete trifurcation, such as in the case of hybrid speciation wherein two topologies significantly exceed the third one but do not significantly differ from each other; (iii) incomplete bifurcation, such as in the case of asymmetric introgression wherein the proportion of all topologies significantly differ; and (iv) complete bifurcation, one topology significantly exceeds the two others, which in their turn have nearly equal proportions. Categories ii and iii are often considered to evidence reticulate evolution. The earliest of introgression tests, Patterson’s *D* or the ABBA-BABA statistic (Green et al. 2010; Durand et al. 2011), compared the two later cases (iii and iv), *i.e.* it presumed that a “true” species tree exists. A later test, HyDe (Blischak et al. 2018), quantifies admixture (γ) from the ratio of shared alleles with the test going from 0 (full isolation) to 0.5 (full hybridization) and therefore it can also cover case ii. The two tests differ in how they measure significance, using jackknifing or bootstrapping in Patterson’s *D* and normal approximation in HyDe. Of late, another site-based test was developed using χ^2^ to test for deviation of parity between the three topologies as in case i (Sayyari and Mirarab 2018). A unified test that can test the prevalence of each of the four categories across the genome is still lacking.

**Figure 1.**
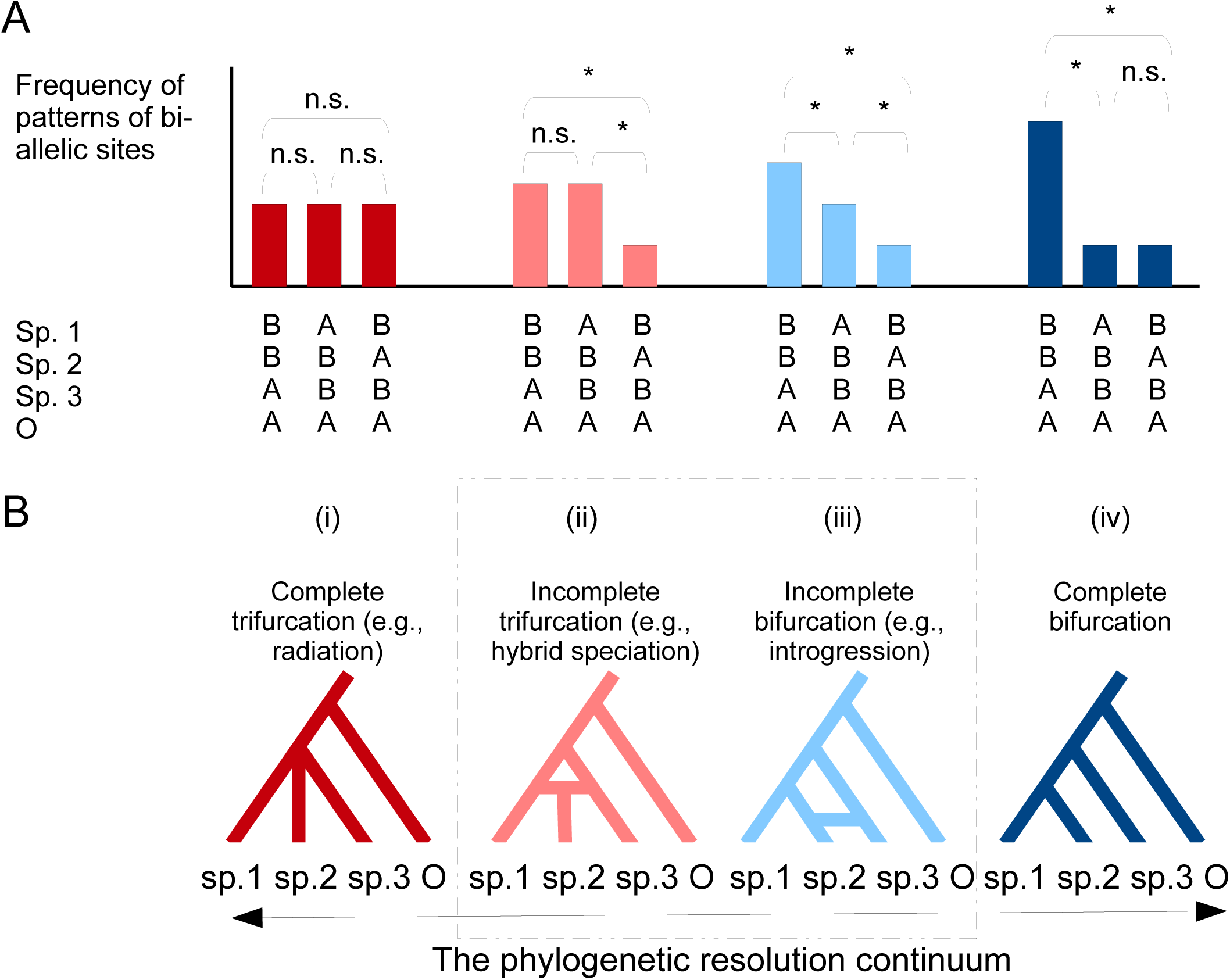
The distribution of bi-allelic sites proportions among three ingroup species with an outgroup and the rationale of the 2A2B test. A) Statistical pairwise comparisons of the three possible bi-allelic patterns generate four categories. B) Each category corresponds to a possible evolutionary scenario along the reticulation-bifurcation continuum. B) Based of the frequency of these topologies in a genome fragment, this fragment can be categorized in (i) complete reticulation, (ii) incomplete reticulation, (iii) incomplete bifurcation, and (iv) complete bifurcation. The 2A2B test estimates significant deviation from parity in pairwise comparisons of the topologies. *: significant deviation, ns = non-significant.

Reminiscent to the problem of soft and hard polytomies, wherein the statistical power of a phylogenetic analysis largely depends on the size of a locus, interpreting reticulation is largely influenced by how a locus is defined. For example, at the species level, an analysis on a whole concatenated alignment of a quartet should have a greater power (*i.e.* a total evidence approach). However, sites are often physically linked and evolutionarily-dependent. Consequently, whole genome analyses may be biased towards genomic regions with more variable sites (Martin et al. 2015). Therefore, phylogenetic analyses are usually conducted at individual loci under the multispecies coalescent model, which was shown to outperform concatenation in multiple groups of animals (Jiang et al. 2020). In theory, a window size of ∼100-kb long is appropriate to overcome the non-independence of sites under realistic recombination rates (Pease and Hahn 2015), but the definition of such windows would largely depend on the taxonomic scale of the analysis. For example, at shallow phylogenetic depths, it is customary to use a pseudo-reference genome approach, whereas reads from multiple species are aligned to a reference genome. Here, the distance from the reference genome species can greatly influence the quality of mapping and subsequent analyses due to potential gene duplications and rearrangements (XXX). Alternatively, single-copy orthologous genes, such as those that have been popularized by the BUSCO program, are used (XXX). However, such gene are presumably under strong selection, with the relaxation of selection in some lineages may conflate introgression estimates (Frankel and Ané 2023). Proper estimation of reticulation therefore requires a good understanding of how different locus-definition strategies influences incongruence analyses.

Phylogenetic conflicts have been reported for the jumping fly *Drosophila saltans* species group, a clade of the subgenus *Sophophora* with 23 Neotropical species (Magalhães 1962; Bächli 2024). The group retains its name from the peculiar “jumping” habit of its larvae; *“the larva seizes its posterior end with its mouthhooks, and stretches. The hooks pull loose suddenly, the larva straightens with considerable force, and as a result is thrown several inches into the air”* (Sturtevant 1942). The group was divided into five species subgroups, namely, *saltans, parasaltans, cordata, elliptica* and *sturtevanti* subgroups, mostly on the basis of male genitalia (Magalhães and Björnberg 1957). Although the monophyly of the subgroups has been confirmed by different phylogenetic methods, the relationships among and within them are not. Hypothesis for their evolutionary relationships have been proposed using different methods and different morphological characters (Magalhães and Björnberg 1957; Throckmorton 1962; Throckmorton and Magalhães 1962; O’Grady et al. 1998; Yassin 2009; Souza et al. 2014; Roman et al. 2022), chromosome polymorphism (Bicudo 1973a), reproductive isolation (Bicudo 1973b; Bicudo and Prioli 1978; Bicudo 1979), protein polymorphism (Nascimento and Bicudo 2002) and gene sequences (Pélandakis and Solignac 1993; O’Grady et al. 1998; Rodrı guez-Trelles et al. 1999; Castro and Carareto 2004; de Setta et al. 2007; Roman et al. 2022). The evolutionary relationships proposed are summarized in Supplementary Table S1.

Unlike other species groups in the subgenus *Sophophora*, such as the *melanogaster*, *obscura* and *willistoni* groups, genomic resources and genetic investigations in the *saltans* species group are scarce. Indeed, only four genomes have been sequenced and assembled to date (Kim et al. 2021). To bridge this gap and to test for the extent of phylogenetic conflicts, we sequenced and assembled genomes for 15 species with representatives from the five subgroups. Phylogenetic analyses using well-conserved genes resolved the evolutionary relationships among the subgroups but also highlighted conflicts between X-linked, autosomal and mitochondrial loci. To test how each of the four incongruence categories prevails across the genome, we devised a new χ^2^-based test that we called 2A2B. The test uses pairwise comparisons of the three topologies proportions shown in Figure 1. We applied this test to sets of conserved orthologous genes and to long syntenic blocks with conserved collinearity across the Neotropical *Sophophora*, as well as to 100-kb long windows from pseudo-reference genomes aligned to either an ingroup (*D. sturtevanti*) or outgroup (*D. willistoni*) species. We found reticulation levels to differ among the datasets and the subgroups, and to correlate with rate of speciation and historical biogeography in the *saltans* group.

## Results

### Short-read assembly of 17 genomes recovered 90% of BUSCO genes

We sequenced using short-read Illumina approach 17 whole genomes from 15 species collected across various locations in the Neotropical region. Genome size, estimated from 21-kmer frequency spectrum using GenomeScope 2 (Ranallo-Benavidez et al. 2020), ranged from 154.0 to 356.8 Mb. Our de novo assemblies using MaSuRCA (Zimin et al. 2013) resulted in genome lengths ranging from 177.5 to 287.7 Mb, with N50 values ranging from 2 to 92 Kb (Supplementary Table S2). To evaluate the completeness of our assembled genomes, we searched for single-copy genes (SCG) using BUSCO (Simão et al. 2015). We found that over 90% of the searched genes were complete for all of the genomes (Supplementary Table S2). Kim et al. (2021) assembled using both short Illumina and long Nanopore reads the genomes of four *saltans* group species, all of which we have independently sequenced. Whereas their assemblies’ contigs were much longer, with N50 ranging from 2 to 6 Mb, the BUSCO score for the same set of species did not largely differ (98% vs. 95-96% in our study; Supplementary Table S2).

### Muller elements analysis resolves relationships between the subgroups and unravels a minor X-autosomal conflict in the sturtevanti subgroup

Phylogenomic analyses were performed using 2,159 SCG shared across all species. Gene trees, inferred for each SCG using maximum-likelihood in IqTree2 (Minh et al. 2020) produced 1,263 distinct topologies, with 206 of them found more than once. To test if SCG chromosomal position may underlie the discrepancies in gene trees, we localized each SCG to its corresponding Muller element according to the position of its *D. melanogaster* ortholog identified by Blast (Camacho et al. 2009). As a result, we generated five independent datasets, each corresponding to the Muller elements A, B, C, D, and E, comprising 337, 370, 425, 419, and 568 SCGs, respectively. These datasets were then used to reconstruct the species trees using the multi-species coalescent model, and the genes within them were concatenated for Bayesian and Maximum Likelihood phylogenetic inferences.

The trees generated by the 5 data sets showed very similar topologies with well supported nodes either in the Bayesian Inference implemented in BEAST (Bouckaert et al. 2019), the maximum-likelihood implemented in IqTree2, or for the multi-species coalescent model analysis implemented in ASTRAL-III (Zhang et al. 2018) (Supplementary Figures S1, S2 and S3). The *parasaltans* subgroup was placed as sister to all other subgroups, followed by the emergence of the *sturtevanti* subgroup. The *cordata* and *elliptica* subgroups showed a close relationship, and were sister to the *saltans* subgroup. The only discrepancy between the topologies was the placement of *D. lehrmanae*, a newly discovered species in the *sturtevanti* subgroup (Madi-Ravazzi et al. 2021). For *D. lehrmanae*, while maximum-likelihood and multi-species coalescent analyses reported lack of branch support for multiple trees (Supplementary Figures S2 and S3), Bayesian inference recovered well supported branches and two topologies (Figure 2A and Supplementary Figure S1). The two distinct topologies distinguished the Muller elements forming the X chromosome (elements A and D who are fused in a fusion shared by the Neotropical *Sophophora*, *i.e.* the *saltans* and *willistoni* groups (Sturtevant and Novitski 1941; Dobzhansky and Pavan 1943; Cavalcanti 1948), and the Muller elements representing autosomal chromosomes (elements B, C and E).

**Figure 2.**
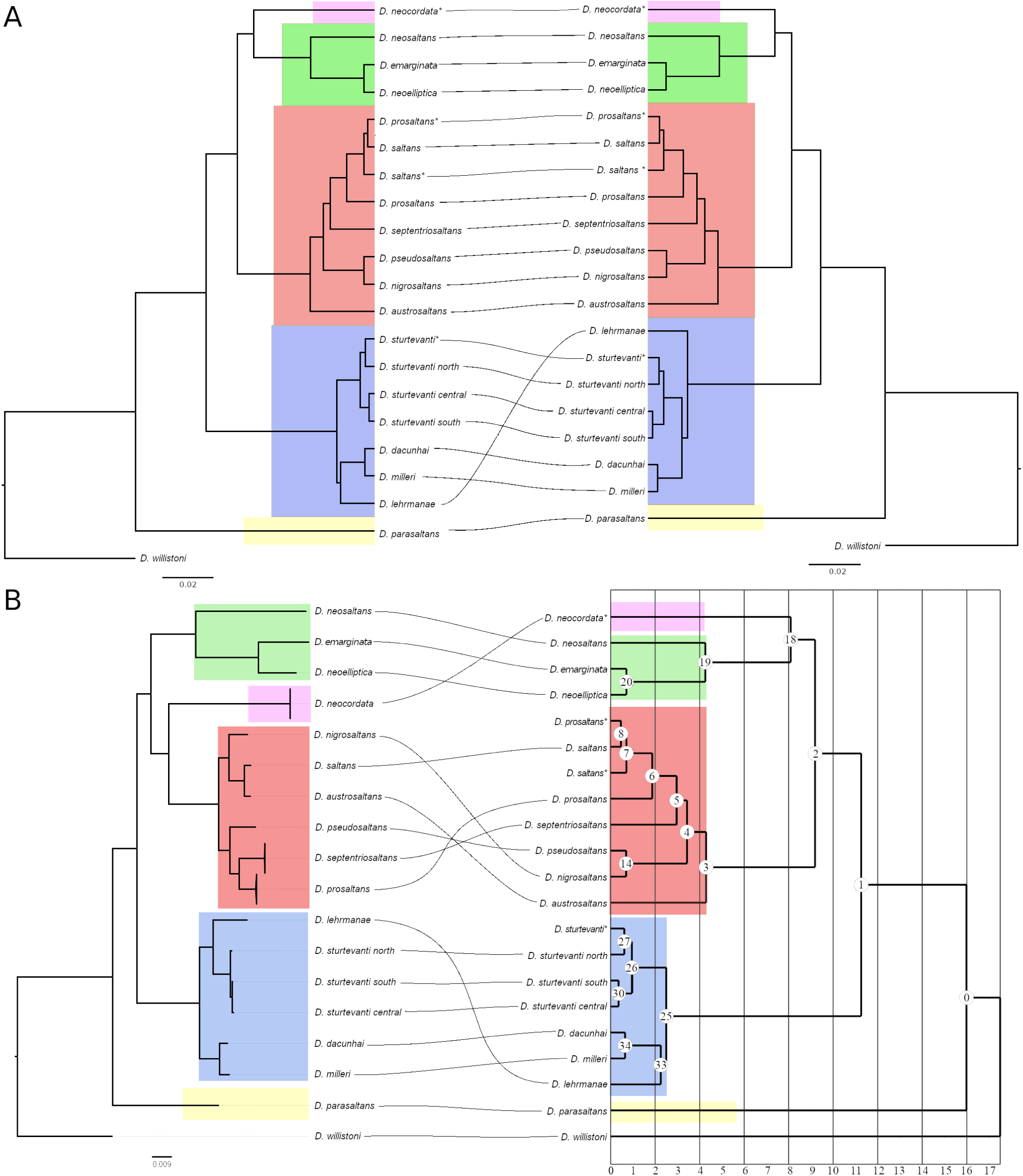
Phylogenomic conflict between the X chromosome, autosomes, and the mitochondria. A) Comparison of autosomal topology (left, represented by Muller element B) and X chromosome topology (right, represented by Muller element A) demonstrates overall agreement with minor incongruence. B) Mitochondrial-nuclear disagreement highlights stronger incongruence between mitochondrial topology (left) and sexual chromosome topology (right). All trees were inferred through Bayesian analysis. Divergence time estimation (in million years ago, myr) for the sexual chromosome topology is provided (left). Nodes are numbered for subsequent analyses. Maximum-likelihood and ASTRAL-III trees are provided in Supplementary Figures S2 and S3, respectively. Posterior probabilities of all nodes were equal to1.

The published genome of *D. prosaltans* (Kim et al. 2021) did not group with the genome of this species sequenced by us, instead it grouped with *D. saltans*. The genome previously published comes from a line collected in El Salvador in 1957. According to Magalhães’ (1962) detailed morphological revision of multiple geographical specimens of the *saltans* group, the sampling site of this particular strain is outside the geographical range of *D. prosaltans*, but within the expected range of *D. saltans*. Furthermore, the *D. saltans* and *D. prosaltans* lines used in our study underwent thorough morphological analyzes (Souza et al. 2014; Roman and Madi-Ravazzi 2021), indicating that the lines we used were accurately identified. Therefore, it is most likely that the previously sequenced *D. prosaltans* strain from El Salvador was misidentified and we consider it here to belong to *D. saltans*.

### Mitogenomes show cytonuclear conflicts in the sturtevanti and saltans subgroups

We assembled mitochondrial genomes for the 15 *saltans* group species using MitoZ (Meng et al. 2019). We did not use the previously assembled four strains since several mitochondrial scaffolds were likely removed in those assemblies (Kim et al. 2021). We conducted phylogenetic analysis on the aligned mitogenomes genes using both IqTree2 and MrBayes (Ronquist et al. 2012). Overall, the mitochondrial trees matched the topology of the nuclear gene trees regarding the inter-subgroup relationships. However, three major discrepancies were identified (Figure 2B). First, the position of *D. lehrmanae* within the *sturtevanti* subgroup did not agree with either the X or autosomal SCG topologies, proposing a topology wherein *D. lehrmanae* is a sister species of *D. sturtevanti* (a topology recovered once in multi-species coalescent analysis (Muller element C, Supplementary Figure S3) and maximum likelihood (Muller element B, Supplementary Figure S2)). Second, whereas the mitochondrial tree recovered the monophyletic relationship between the *elliptica*, *cordata* and *saltans* subgroups, the position of *D. neocordata* (*cordata* subgroup) differed, being sister to the three species of the *elliptica* subgroup in the nuclear trees and to the six species of the *saltans* subgroup in the mitochondrial tree. Third, whereas nuclear trees recovered three lineages within the *saltans* subgroup, namely*, austrosaltans*, *nigrosaltans-pseudosaltans*, and *septentriosaltans-prosaltans-saltans*, only two lineages are revealed by the mitochondrial tree. Each of the mitochondrial clades involved one species from otherwise sister species in the nuclear trees, *i.e. D. nigrosaltans* and *D. saltans* in one clade and their respective closely-related species *D. pseudosaltans* and *D. prosaltans* in the other clade, suggesting that multiple cytoplasmic introgression events might have occurred in this subgroup (Figure 2B). Because *D. saltans* and *D. prosaltans* are reported as closely related species in the nuclear trees and are separated in the two mitochondrial clades, the two mitoclades were called *S* and *P*, respectively. Remarkably, the branching order in the *P* mitoclade is congruent with the nuclear tree, indicating that the *S* mitoclade is most likely of reticulate origin, probably reflecting an older introgression from *D. austrosaltans* to *D. nigrosaltans* and a more recent introgression between *D. austrosaltans* and *D. saltans*.

### The extent of reticulation differs between assembly-and pseudo-reference-based approaches

To quantify the extent of introgression at a genome-wide scale, we analyzed allele distribution of bi-allelic phylogenetically informative sites in 28 species quartets using Patterson’s *D* estimate of the standard ABBA-BABA test. We used four distinct locus-defining strategies to explore how strategy choice can influence estimates of *D*. Two strategies relied on our *de novo* genome assemblies. The first strategy included the use of single-copy genes (SCG) defined by BUSCO (2,159 loci). The second strategy identified syntenic blocks equal to or greater than 50 kb-long with collinearity across the 15 *saltans* assemblies and *D. willistoni*, *i.e.* across Neotropical *Sophophora* (1,797 loci). The third and fourth strategies used a pseudo-reference approach, wherein the reads of each *saltans* group species were mapped to an ingroup (*D. sturtevanti*, 2,328 loci) or an outgroup species (*D. willistoni*, 1,863 loci) and a pseudo-reference was inferred by replacing variant sites in the reference genome by the ones in the reads (as in Mai et al. 2020). In these pseudo-reference strategies, a locus was defined as a 100 kb-long window of the reference genome. To avoid the influence of linked loci from highly-variable regions on genome-wide estimates, we randomly sampled 20 sites from each locus with ≥20 evaluated sites, and repeated resampling for 100 reiterations for each locus-defining strategy.

For a bi-allelic site, the standard ABBA-BABA (Patterson’s *D*) test considers a true species tree exists, with site distribution AABB, and then evaluate the deviation from parity of the two discordant configurations ABBA and BABA. We found that genome-wide absolute Patterson’s *D* estimates, *i.e.* (∑ABBA – ∑BABA)/(∑ABBA + ∑BABA), greatly differed among the four strategies (Figure 3A). Synteny-based strategy had the lowest |*D*| across the 28 quartets (|*D*| = 0.025 ± 0.004) compared to BUSCO (|*D*| = 0.066 ± 0.014, Student’s *t P* < 0.01), *D. sturtevanti* pseudo-reference (|*D*| = 0.176 ± 0.025, Student’s *t P* < 4.77 x 10^-6^), and *D. willistoni* pseudo-reference (|*D*| = 0.329 ± 0.053, Student’s *t P* < 5.97 x 10^-6^). So, of the four strategies, the use of syntenic blocks is the most conservative.

**Figure 3.**
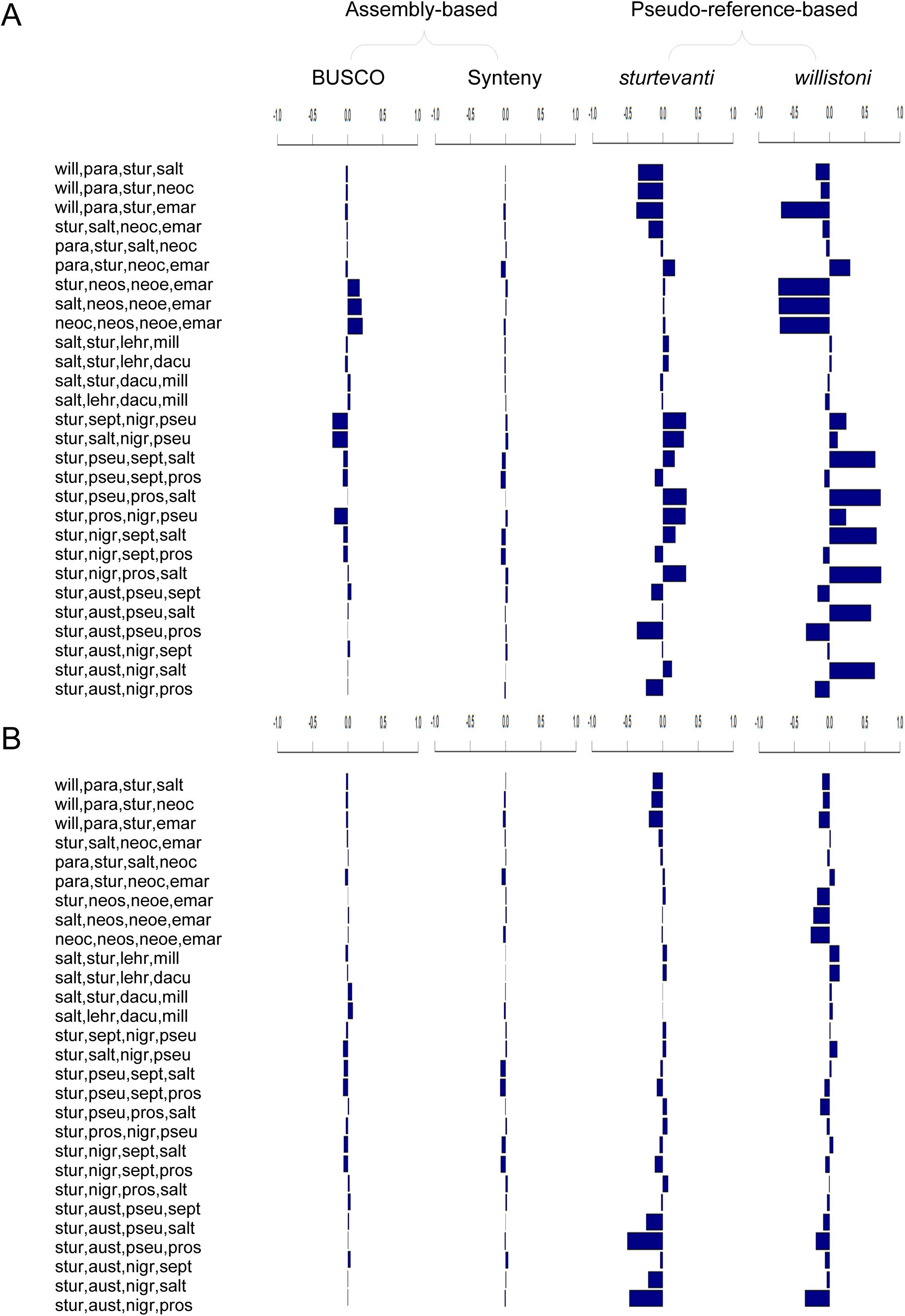
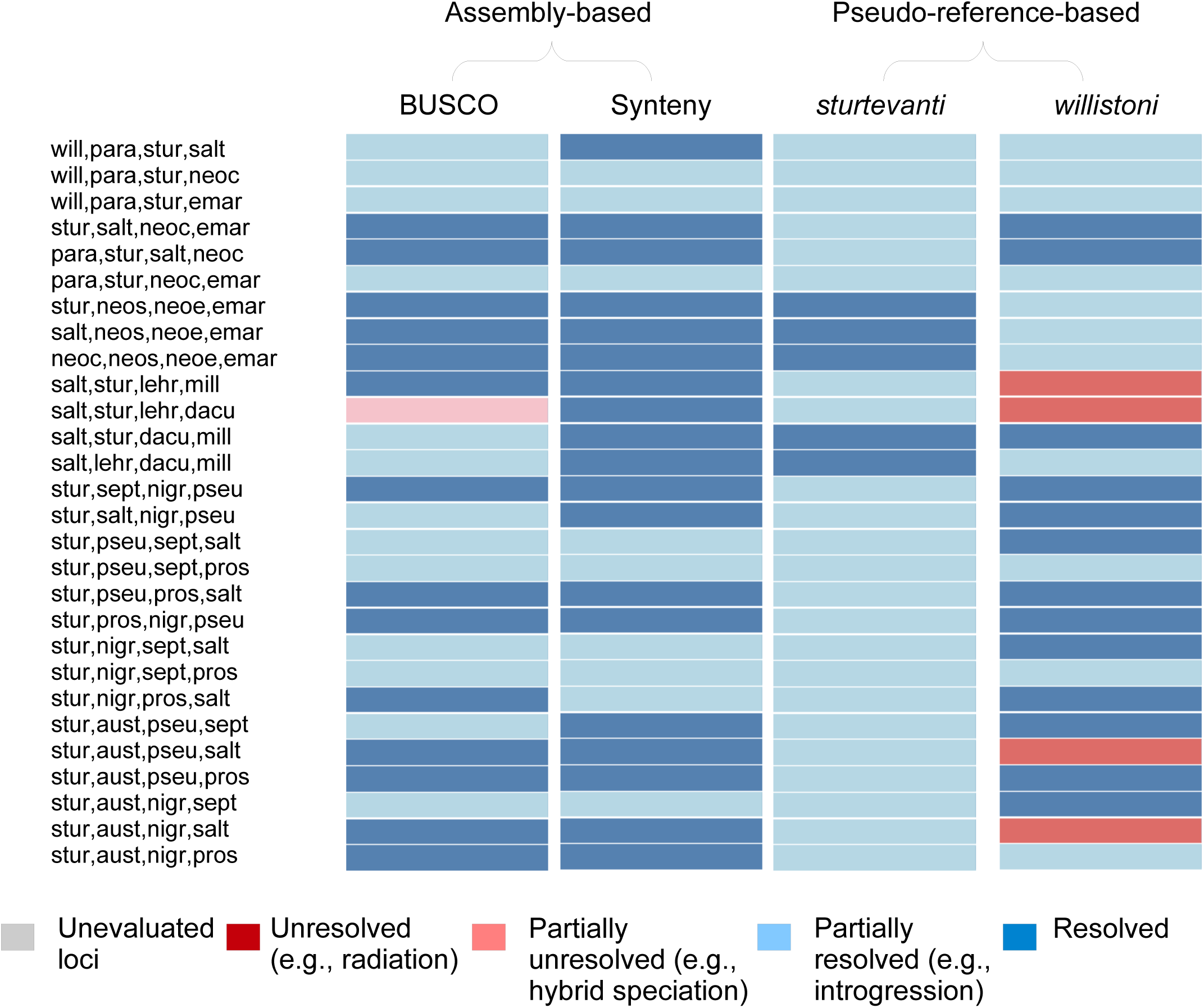
Extent of phylogenomic discordance across 28 species quartets as measured by A) Patterson’s *D* and B) the 2A2B test, using four locus-definition strategies (see text). For both tests, assembly-based strategies have lower average *D* and provide stronger evidence for reticulation at three episodes. Colors of bars in B) correspond to the four categories presented in Figure 1, with grey bars corresponding to loci with less than 20 evaluated bi-allelic sites.

### Comparisons of locus-defining strategies identify sources of errors in whole-genome Patterson’s D estimates of introgression in species quartets

To gain further insights on the reasons of this discrepancy between the four locus-defining strategies, we separately investigated assembly-and pseudoreference-based approaches. The BUSCO approach mainly differs from the synteny-based approach in having, on average, shorter sequence since many loci are ≤50-kb long. Short sequences usually shows faster lineage-specific evolutionary rates (XXX), which can increase variance and also induce biases on Patterson’s *D* due to the violation of the molecular clock assumption (Frankel and Ané 2023). Besides, the BUSCO analysis involves an additional step, which is the annotation of the conserved protein-coding BUSCO genes. To account for these possible sources of errors, we counted for each lineage the number of singletons distinguishing the outgroup from the three ingroup species (*i.e.* BBBA sites), as well as the singletons distinguishing between the two most closely-related species in the quartet as defined from the phylogenetic analyses above (*i.e.* BAAA and ABAA). we called this test 3A1B, and it is reminiscent to the classical Tajima’s (1993) Relative Rate Test. Under the assumption of a constant molecular clock or the absence of mis-annotation of a BUSCO gene in one of the two sister species both BBBA – BAAA and BBBA – ABAA should be > 0. We focused on the *elliptica* subgroup with *D. saltans* as an outgroup (*i.e.* (((*emarginata*,*neoelliptica*),*neosaltans*),*saltans*)) quartet, since this quartet showed a strong discrepancy between the two assembly-based approaches, with genome-wide *D* being 0.25 for BUSCO and -0.08 for synteny-based strategy. By applying the 3A1B test, we found that 37 BUSCO loci violated its assumption, all in the negative direction for *D. emarginata*, *i.e.* BAAA > AAAB. By excluding these loci, Patterson’s *D* dropped from 0.25 to 0.04. By contrast, only 5 loci violated the assumption in the synteny-based strategy. Their exclusion did not change Patterson’s *D* estimate which remained -0.08. We then applied the 3A1B test for all locus-defining approaches, excluding loci that violated its basic assumption in each quartet. The genome-wide absolute Patterson’s *D* estimates dropped from 0.066 ± 0.014 to 0.030 ± 0.004 for BUSCO genes (Student’s *t P* < 0.05) and from 0.025 ± 0.004 to only 0.023 ± 0.004 for synteny-based loci (Student’s *t P* = 0.76). After correction, genome-wide estimates from both BUSCO and synteny loci did not significantly differ (Student’s *t P* = 0.28), even if synteny estimates remained slightly lower (Figure 3B).

For pseudo-reference approaches, excluding rapidly evolving loci through the 3A1B test may not be enough, since additional sources of errors related to the quality and depth of mapping as well as reference genome biases constitute additional sources of errors. We therefore reconducted the analyses using more stringent parameters, only retaining loci with a mapping quality >20 and a depth >10 and <100 and excluding all within-species polymorphic sites. These parameters dramatically reduced the number of evaluated loci to only 17% and 4% in the ingroup (*D. sturtevanti*) and outgroup (*D. willistoni*) reference genome approaches. Estimation of genome-wide absolute Patterson’s *D* to 0.101 ± 0.023 (Student’s *t P* < 0.05) and 0.102 ± 0.017 (Student’s *t P* < 2.9 x 10^-4^) for the ingroup and outgroup references, respectively. This means that whereas ingroup and outgroup approaches were greatly different before the application of the stringent criteria (Student’s *t P* < 0.05), absolute Patterson’s *D* did not significantly differ between the two approaches under these criteria (Student’s *t P* = 0.95). For the abovementioned *elliptica* subgroup example, whereas before the stringent criteria Patterson’s *D* were 0.02 and -0.71 for ingroup and outgroup approaches, these values dropped to -0.005 and -0.27 in the two approaches, respectively. However, correlation across quartets was strong (Pearson’s *r* = 0.56, *P* = 0.002), indicating that distinct criteria may be applied to improve estimate depending on the distance from the reference genome species, with *D. willistoni* being more distant and having a more fragmented genome assembly.

### The 2A2B test unravels lower signal for introgression when syntenic blocks are used

To consider cases where a true species tree cannot be inferred, we designed a test that we call 2A2B. The test is an extension of the standard ABBA-BABA test in that it does not only test the significance of deviation from parity of the ABBA and BABA patterns, but also test the deviation from parity of each configuration to the BBAA pattern. Therefore, the test allows classifying each species quartet into one of the four categories along the trifurcation-bifurcation continuum given in Figure 1. We applied the test at both the genome-wide level after corrections using the 3A1B test and mapping criteria (see above).

We found that for all locus-defining strategies, bifurcation categories (iii and iv) predominated (Figure 4A). Only five exceptions were observed. One incidence of category ii (incomplete trifurcation) was observed in the BUSCO strategy concerning the position of *D. lhermanae* relative to both *D. sturtevanti* and *D. dacunhai*. Indeed, the position of *D. lhermanae* was the only conflicting result in the phylogenetic analyses based on BUSCO genes and it also showed a conflict with mitochondrial analysis (Figure 2). Four incidences of category i (complete trifurcation) were observed in outgroup pseudo-reference strategy. Of which, two concerned the position of *D. lehrmanae* to both *D. sturtevanti* on the one hand and to each of the two sister species *D. dacunhai* and *D. milleri* on the other hand. The other two complete trifurcation cases concerned the relationships between the three lineages of the *saltans* subgroups, *i.e. austrosaltans*, *nigrosaltans-pseudosaltans*, *septentriosaltans-prosaltans-saltans*, in cases where species of the different mitoclades where found analyzed together, *i.e. D. pseudosaltans* and *D. saltans* or *D. nigrosaltans* and *D. septentriosaltans* (Figure 2).

**Figure 4.**
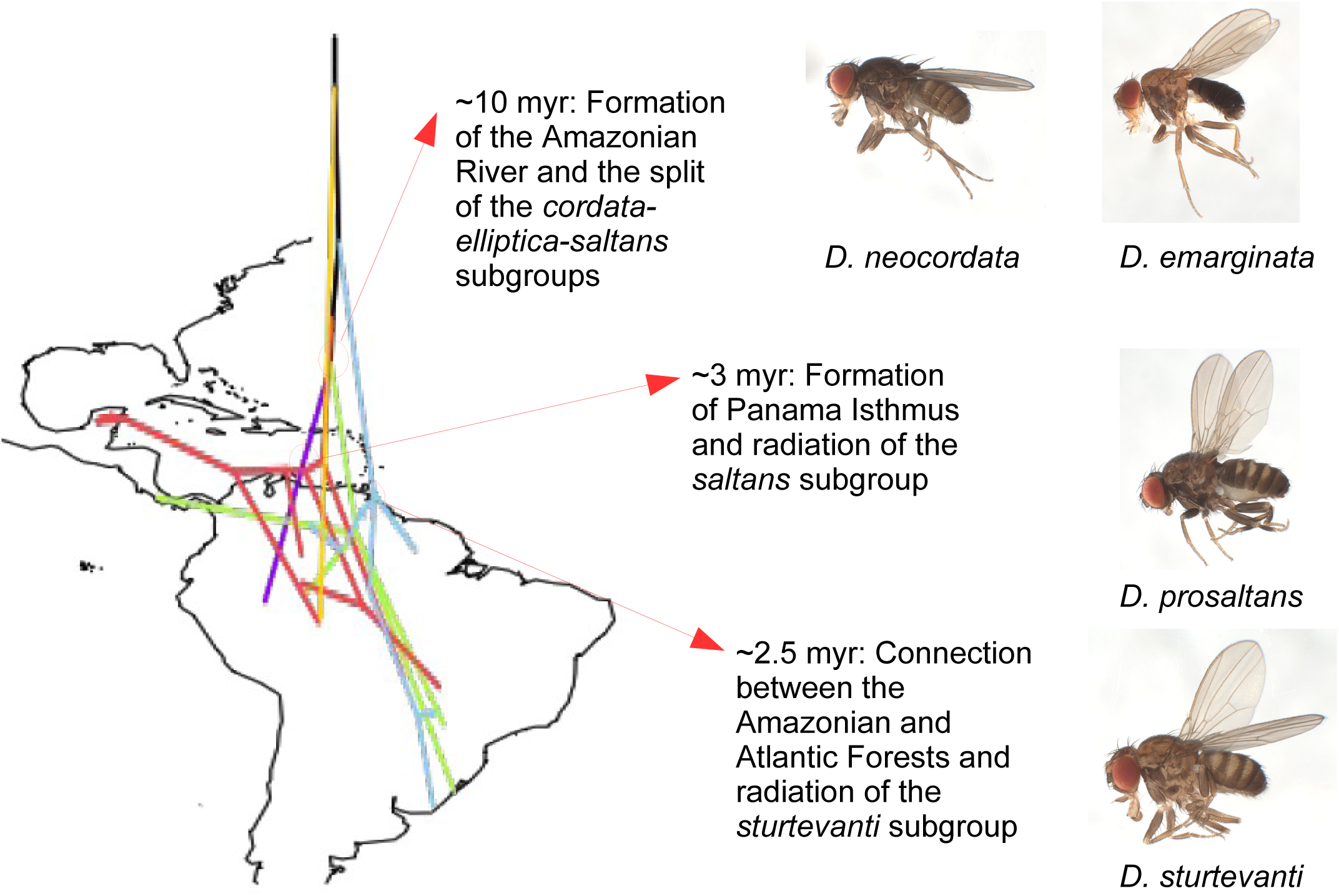
Historical biogeography of the *saltans* species. Centroid geographical coordinates for each species were inferred and ancestral coordinates were inferred using BayesTraits on the Bayesian X chromosome chronogram given in Figure 2B. Internal nodes are projected on the current map of South and Central America. Three episodes of rapid radiation were identified by the 2A2B test (Figure 3B) and are identified here with their corresponding possible paleogeographical events, with photos of adult males of four representative species shown. Branch colors in the tree correspond to the subgroups colors given in Figure 2.

Category iv (complete bifurcation), which is the most consistent with a true species tree, was the most frequent in synteny-based strategy (Figure 4). This was followed by both the BUSCO and the outgroup pseudo-reference strategies but the difference in category iv frequency was not significant (Fisher’s exact test of category iv vs. all other categories). However, with only four exceptions, ingroup pseudo-reference approach identified category iii (incomplete bifurcation), which mostly indicates cases of introgression, in every possible quartet on the tree (Fisher’s exact test between synteny-based and ingroup pseudo-reference strategy *P* < 1 x 10^-4^). The total number of bi-allelic sites (20 per locus) that were used in the genome-wide analysis differed among the four strategies, averaging 22,893 (BUSCO) and 21,306 (synteny) assembly-based approaches, and 35,747 and 5,308 sites for the ingroup *D. sturtevanti* and the outgroup *D. willistoni* pseudo-reference approaches, respectively. The low number for the *D. willistoni* pseudo-reference is expected since increasing mapping quality criteria on the genome of this distant species would only favor strongly conserved loci. For the *D. sturtevanti* pseudo-reference, applying the same mapping quality criteria has likely maintained less conserved loci. Although we have excluded heterozygous sites from every species to reduce mapping biases of the reference allele, such biases may even extend to homozygous sites linked to the excluded heterozygous site in the same read. Universal mapping criteria that can reduce mapping biases independent from the reference genome do not exist and mapping biases remain difficult to detect (XXX). However, these biases are the most likely explanation of the extensive signal of introgression in the ingroup *D. sturtevanti* pseudo-reference. To test the effect of mapping quality on introgression estimates, we applied the 2A2B test at the locus level before and after applying mapping quality thresholds. In agreement with our assumptions and the findings on Patterson’s *D* (Figure 3), the proportion of loci supporting categories ii and iii, *i.e.* categories that are the most associated with reticulate evolution, did not exceed ∼7% in assembly-based strategies, but they dropped after application of mapping quality criteria from 46% to 11% and from 34% to 4% for *D. sturtevanti* and *D. willistoni* pseudo-reference genomes strategies, respectively (Supplementary Figure XXX). Ingroup pseudo-reference approaches, while providing more data, may not reflect the true phylogenetic relationships between the species, hence inflating the estimation of the extent of introgression.

### Historical biogeographical events coincided with episodes of low phylogenetic resolution

Phylogenetic analyses and assembly-based 2A2B test identified three parts on the species tree with evidence for incomplete bifurcation (*i.e.* category iii): at the inter-subgroup level in any combination that involved representatives from at least two subgroups of the *cordata*, *elliptica* and *saltans* subgroups, at the base of the *sturtevanti* subgroup, and in the *saltans* subgroup in quartets involving species belonging to distinct mitoclades. Whereas category iii is usually interpreted as evidence of introgression following secondary contact, other biological processes such as ancestral population structure at the time of phyletic radiation or asymmetry of evolutionary rates due to responses to distinct adaptive pressures (XXX). Because all these processes can associate with major geographical changes affecting species distribution, ecology and frequency of secondary contacts, a knowledge of the historical biogeography of the clade under study is crucial for the understanding of the processes that have likely led to phylogenetic conflicts.

Like phenotypic traits, ancestral geographical distribution can be inferred on phylogenetic trees from present day geographical data (XXX). Therefore, we mapped current distribution of the studied species on the Bayesian X tree that was dated in reference to recent family-wide estimates (Suvorov et al . 2021; see Methods). The historical biogeography supported an origin of the *saltans* group around 16 million years (myr) ago in the north-west of South America, which currently corresponds to the Amazonian region (Figure 5). At that time, this region was dominated by a vast wetland environment known as the ‘Pebas System.’ The first radiation episode, *i.e.* the split between the ancestors of the *cordata*, *elliptica* and *saltans* subgroups occurred circa 10 myr ago at this region, coinciding with the formation of the Amazon River and its huge fluvial networks that led to the transformation of the Pebas wetland into the dense and diverse Amazonian forests (Hoorn et al. 2022). The second episode occurred nearly 3 myr ago, which correlates with the geological formation of the isthmus of Panama (O’Dea et al. 2016), and the time of a connection between the Western Amazonian and the Southern Atlantic rainforests during the Pliocene (Batalha-Filho et al. 2013; Ledo and Colli 2017). This episode involved the ancestors of the *nigrosaltans-pseudosaltans* and *septentriosaltans-prosaltans-saltans* clades. Indeed, the first clade has a northern to central representative (*D. nigrosaltans*) and a southern species (*D. pseudosaltans*), whereas the second clade has a northern representative (*D. saltans*) and two central and central to southern species (*D. septentriosaltans* and *D. prosaltans*, respectively). The third radiation episode involves the ancestor of the *sturtevanti* subgroup and also occurred in the late Pliocene, ca. 2.5 myr ago, at the likely time of a connection between the Western Amazonian and the Southern Atlantic rainforests.

## Discussion

### Towards a comprehensive phylogeny of the saltans species group

A large number of *Drosophila* genomes have been sequenced and used in phylogenetic analyses (Khallaf et al. 2021; Kim et al. 2021; Li et al. 2022; Suvorov et al. 2022), but studies with comprehensive sampling of nearly all species in a group remain relatively uncommon (Mai et al. 2020; Conner et al. 2021; Yusuf et al. 2022; Moreyra et al. 2023). Despite minor inconsistencies, our phylogenomic analysis of 15 species of the *Drosophila saltans* species group produced a consistent picture of the relationships between the five subgroups of this clade. All X, autosomal and mitochondrial phylogenies showed the *parasaltans* subgroup as the first to diverge, followed by the *sturtevanti* subgroup, and later by a clade comprising the *cordata*, *elliptica* and *saltans* subgroups, in which the position of the *cordata* subgroup differed between nuclear and mitochondrial trees. This general picture has not been previously proposed despite the tremendous number of phylogenetic investigations of this group (Magalhães 1962; Throckmorton 1962; Throckmorton and Magalhães 1962; O’Grady et al. 1998; Rodrı guez-Trelles et al. 1999; Castro and Carareto 2004; de Setta et al. 2007; Yassin 2009; Souza et al. 2014; Roman et al. 2022; see Supplementary Table S1).

While our analysis has shed light on the intricate evolutionary dynamics within this clade, further sampling holds the potential to provide a more comprehensive understanding into this complex evolutionary history. For example, the inclusion of *D. subsaltans*, *D. lusaltans*, *D. cordata*, and *D. rectangularis* through whole-genome sequencing promises to provide insight onto unresolved phylogenetic questions raised from previously published observations on reproductive isolation and morphology (Magalhães 1962; Bicudo and Prioli 1978). These questions include the monophyly and positioning of the *parasaltans* and *cordata* subgroups. Additionally, the inclusion of the insular species *D. lusaltans* which presents low reproductive isolation (Bicudo 1973b), can bring new insights into the reticulation evolution in the *saltans* subgroup. The *saltans* subgroup showed the most dramatic signal of cyto-nuclear discordance and reticulated evolution. Bicudo (1973b) investigated reproductive isolation among the seven then described species of this subgroup, and in a remarkably partial agreement with our nuclear phylogenomic trees, she concluded that *D. pseudosaltans*, *D. nigrosaltans* and *D. austrosaltans* showed more basal relationships compared to *D. lusaltans*, *D. septentriosaltans*, *D. prosaltans* and *D. saltans*, *i.e.* the latter species show less reproductive isolation. Indeed, nearly all crosses among the last four species produce fertile females with some even producing fertile females and males (Bicudo 1973b). This behavioral porosity largely agrees with the high incidence of reticulate evolution we report here for this subgroup.

Two widespread species of the *saltans* subgroup, *D. saltans* and *D. prosaltans*, show a peculiar geographical disjunction, with the former species being restricted to Central America whereas *D. prosaltans* is widespread in South America south of Costa Rica. The discrimination between strains belonging to each species has long been erroneous (Dobzhansky 1944; Mayr and Dobzhansky 1945; Spassky 1957; Magalhães 1962) and we showed here that their misidentification persists even in the genomic era (Kim et al. 2021; Suvorov et al. 2022). Interestingly, Bicudo (1973b) provided evidence for reproductive reinforcement between these two sister species; sympatric populations in their junction zone in Costa Rica demonstrated stronger reproductive isolation than allopatric populations of both species. We have only included one to a few geographical lines from each species and a broader sampling to investigate the extent of their reproductive isolation and genome porosity is strongly needed.

### Intra-and inter-genomic conflicts impact the inference of phylogenetic patterns

Concatenation helped recovering a sex chromosome versus autosome conflict, similar to the one described by Mai et al. (2020) for the *Drosophila nasuta* subgroup. Like these authors, this conflict was limited to a single part of the tree, *i.e.* the relationship of *D. pulau* to *D. sulfurigaster sulfuricaster* and *D. s. bilimbata* in the *nasuta* group and the placement of *D. lehrmanae* in the *sturtevanti* subgroup. The peculiarities of sexual chromosomes such as their lower effective population size, different recombination and mutation rates, and greater exposition to natural selection when found in hemizygosity, lead to higher rates of adaptive evolution of sexual-linked genes compared with autosomal genes (*i.e.* faster-X evolution) and also to the disproportional accumulation of genes related to reproductive isolation and Dobzhansky-Muller hybrid incompatibilities (*i.e.* Haldane’s rule) (Charlesworth et al. 2018). Besides, low recombination rate underlies stronger linked selection effects against introgressed ancestry. Altogether, those characteristics are thought to be responsible for the resistance to hybridization in the sexual chromosomes (Ellegren 2009; Qvarnström and Bailey 2009; Sankararaman et al. 2016; Charlesworth et al. 2018; Seixas et al. 2018; Mai et al. 2020; Matute et al. 2020; Moran et al. 2021; Reilly et al. 2022; Skov et al. 2023; but see David et al. 2022).

The second conflict regards a significant disagreement between mitochondrial (mtDNA) and nuclear data. Discordance between nuclear and mitochondrial genomes is a well-documented phenomenon in the tree of life as highlighted by Toews and Brelsford (2012). Several characteristics of mtDNA, such as being haploid and uniparentally inherited, resulting in a fourfold reduction in effective population size when compared with autosomal chromosome loci, affect its evolution. Cytoplasmic introgression has long been recognized in *Drosophila* (Solignac et al. 1986; Ballard 2000; Llopart et al. 2014). In a recent population study within the *willistoni* group, multiple mitochondrial introgressions were observed in *D. paulistorum* populations (Baião et al. 2023). These included an ancient introgression with a highly divergent mitochondrial type, followed by more recent events. While nuclear-mitochondrial incompatibilities likely posed challenges, the study also suggested two possible alternatives to overcome these challenges: a selective advantage provided by the mitochondrial type itself. Or a non-selective factor, such as *Wolbachia,* a bacteria known to modify the reproduction of its host, could facilitate a mitochondrial type fixation (Baião et al. 2023). Although, interesting results have been reported from population approaches, conflicts between nuclear and mitochondrial genomes have not been addressed in recent phylogenomic analyses in the Drosophilidae (Mai et al. 2020; Khallaf et al. 2021; Suvorov et al. 2022; Yusuf et al. 2022). The disagreement was particularly evident for the *saltans* subgroup, where it was most likely of recent origins, separating species that have diverged only 0.7 myr ago, *i.e. D. nigrosaltans* and *D. pseudosaltans*. Remarkably, the two mitotypes P and S do not correlate with the degree of reproductive isolation inferred by Bicudo (1973b), contrary to nuclear tree, indicating that cytoplasmic introgression in the *saltans* subgroup did not contribute to the evolution of reproductive isolation in this clade.

Of the three subgroups for which multiple species were sequenced, the *saltans* subgroup had the highest incidence of incongruences inferred by the 2A2B test. In this subgroup, multiple large chromosomal inversions are known to be shared among closely-related species (Dobzhansky and Pavan 1943; Cavalcanti 1948; Bicudo 1973a; Bicudo and Prioli 1978) and evidence for balancing on ancestral inversion polymorphism has been demonstrated in a number of cases (Bicudo 1973a). For example, one autosomal inversion is polymorphic in *D. austrosaltans*, with one arrangement being homozygous in *D. septentriosaltans* and *prosaltans*, and the alternative arrangement homozygous in the remaining species including *D. saltans*. Such an inversion has likely been inherited from a polymorphic ancestor of the *saltans* subgroup, and maintained polymorphic in the ancestor of the *septentriosaltans-prosaltans-saltans* clade before different arrangements became fixed in each of these species, with one arrangement being already fixed in the ancestor of the *nigrosaltans-pseudosaltans* clade. Alternatively, the inversion might have originated in the ancestor of the *septentriosaltans-prosaltans-saltans* clade, then it was introgressed in *D. austrosaltans*, whereas the ancestral non-inverted configuration being introgressed on its own in *D. saltans*. However, this introgression scenario is less parsimonious, especially given the geographical isolation of *D. saltans* from the remaining species of the subgroup. Whether the high degree of incongruences in the *saltans* subgroup is associated with large ancestral inversions potentially absent in other bifurcating clades would require the future generation of chromosome-level assemblies for multiple *saltans* group species.

### The 2A2B test on syntenic blocks helps distinguishing “soft” and “hard” introgressions

Our analyses reveal that the choice of locus defining scheme can affect the degree of phylogenomic uncertainty. Indeed, most studies that have provided strong evidence for introgression using site-specific patterns comparisons have usually aligned reads from multiple species to a single reference assembly (cf. studies compiled by Dagilis et al. 2021). Whereas such an approach would increase the power (Martin et al. 2015; Pease and Hahn 2015), it also introduces biases due to paralogy, misalignments or absence of collinearity among species, that are now known to bias phylogenetic inference itself (Valiente-Mullor et al. 2021; Rick et al. 2023). Therefore, the most common practice now is to use sets of clade-specific single-copy orthologs, such as the BUSCO genes, in inferring phylogenies (Manni et al. 2021). Consequently, some recent introgression studies have focused on such genes. For example, in a study of 155 genomes covering a wide range of drosophilid lineages, Suvorov et al. (2022) analyzed these conserved genes. While finding evidence for widespread introgression, they also showed that their discordance estimates were highly sensitive to the length of the analyzed genes as well as by the slightest relaxation of selective pressures. We attempted here to overcome this limitation by including larger syntenic blocks, which in addition to conserved protein-coding sequences should also include presumably neutral intronic and intergenic sites. Whereas, this approach showed the lowest |*D*|, indicating that patterns predicted by introgression do not exceed 7% of the genome, it led to only 1.5-fold increase in the number of analyzed sites, most likely because our assemblies were based on only short reads. With the continuing progress in whole-chromosome sequencing techniques and assembly-to-assembly alignment tools, a better understanding of the extent of probably true or “hard” introgression events from artificial or “soft” ones may be attained.

Phyletic radiations occur when isolation barriers simultaneously and rapidly evolve between multiple populations of a species with a broad geographical or ecological breadth. Consequently, genetic lineages will be incompletely sorted in the emerging species leading to a lack of phylogenetic signals. This situation usually occurs following a rapid environmental change or expansion into a new range with new empty niches. Indeed, our historical biogeography approach shows that such situation might have favored the episodic signals of low phylogenetic signals in the *saltans* group since the episodes coincided with either the transition from wetlands into Amazonia forests in the Late Miocene or the connection of the Amazonian forest with either Central American or the Southern Atlantic forests in the Pliocene. These events have been proposed to be responsible for a wide range of rapid radiations in Neotropical taxa, such as butterflies, amphibians and birds (Batalha-Filho et al. 2013; O’Dea et al. 2016; Ledo and Colli 2017; Hoorn et al. 2022). The dating and the analyses of the extent of phylogenetic discordance in the genomes of such clades are strongly needed to identify the impact of these events on Neotropical diversification.

## Materials and Methods

### Sample collection, whole genome sequencing and assembly

We performed whole genome pool sequencing on female flies from 15 different species from the *saltans* group, as well as three populations of *D. sturtevanti*. The specimens used for sequencing were obtained from one or multiple strains, and detailed information regarding the number of individuals and their collection locations can be found in Supplementary Table S2. For all species, DNA extraction, PE library preparation using NEBNext Protocol, and 75x coverage double-paired Illumina Novoseq sequencing were conducted by End2End Genomics LLC, Davis, California. For all species except *D. neocordata*, we assembled the genome using the Maryland Super Read Cabog Assembler (MaSuRCA) (Zimin et al. 2013), which utilizes both the de Bruijn graph and overlap-layout-consensus (OLC) methods to generate super-reads. K-mer based genome size was estimated using KMC3 (Kokot et al. 2017) and GenomeScope v.2.0 (Ranallo-Benavidez et al. 2020). To assess the assembly’s completeness, we searched for single-copy-genes (SCG) using default parameters in Busco5 (Simão et al. 2015) with the diptera_odb10 database (Kuznetsov et al. 2023). In addition to the sequenced flies, we also assembled Illumina reads the genomes of *D. saltans*, *D. neocordata*, *D. prosaltans* and *D. sturtevanti* published by Kim et al. (2021) (assembly numbers ASM1890357v1, ASM1890361v1, ASM1815127v1 and ASM1815037v1, respectively), and we used the *D. neocordata* assembly published by Baião et al. (2023) in all analyses.

### Phylogenomics: Nuclear single-copy-genes

For phylogenomics analysis, SCG searches were carried out using 3,285 SCG from diptera_odb10 database on Busco5 (Simão et al. 2015). SCG present in all species were kept and aligned using the L-INS-i method implemented on MAFFT (Katoh and Standley 2013) (mafft --localpair --maxiterate 1000 –adjustdirection).

Genomic data of different species of *Drosophila* support that genes tend to be situated within the same Muller element across multiple species (see Schaeffer 2018). Taking this gene linkage into account, we reconstructed five independent datasets (Muller elements A-E), each comprising all SCG found in the respective Muller element. To achieve this, we performed a tBlastn search against the *D. melanogaster*, and subsequently we concatenated genes found within the same Muller element. Phylogenetic trees were then constructed using maximum likelihood and Bayesian methods implemented in the softwares IQ-TREE (Nguyen et al. 2015) and BEAST (Bouckaert et al. 2019), respectively. Additionally, maximum-likelihood trees were generated for each gene, the output tree from each Muller element dataset were used to reconstruct to species trees, using multi-species coalesce model implemented in ASTRAL-III (Zhang et al. 2018). Branch support was assigned using 1,000 ultrafast bootstrap in IQ-TREE (Hoang et al. 2018), posterior probability in BEAST and local posterior probability in ASTRAL-III (Zhang et al. 2018).

### Phylogenomics: Mitochondrial Genome

Mitochondrial genomes were assembled and annotated with MitoZ (Meng et al. 2019), with the Megahit assembler (Li et al. 2015). In order to ensure the exclusion of nuclear-embedded mitochondrial DNA sequences within the assembly, a strategic approach was taken. Considering that mitochondrial reads are found in higher frequency than nuclear-mitochondrial DNA sequences, the read subsampling were set to 0.5 gigabases (--data_size_for_mt_assembly 0.5). The genes obtained from the mitochondrial genome were aligned using the MAFFT alignment tool with the --auto parameter due to the close similarity between sequences. Subsequently, the aligned genes were concatenated into a dataset for phylogenetic analysis. The concatenated dataset served as the basis for reconstructing phylogenetic trees using both Maximum Likelihood (ML) implemented in IQ-TREE (Nguyen et al. 2015) and Bayesian Inference (BI) in BEAST (Bouckaert et al. 2019). Branch support was assigned using 1,000 ultrafast bootstrap in IQ-TREE (Hoang et al. 2018) and posterior probability in BEAST.

### Identification of syntenic blocks

The 20 genomes of the *saltans* group, the 15 sequenced here and the 5 published by Kim et al. (2021) and Baião et al. (2023) were preliminary annotated with Miniprot (miniprot -Iut16, Li 2023). The primary objective of this annotation was to accurately map proteins from the robust and reliable genome annotation of *D. willistoni* (Clark et al. 2007). After the protein mapping, the predicted gene loci were assessed to identify syntenic blocks present in the Neotropical *Sophophora* (comprising *D. willistoni* and *saltans* group). The identification of these blocks was based on gene order and orientation, achieved using an in-house Perl script. First, this script compares the scaffolds’ genes order and orientation between the reference assemblies of *D. saltans* and *D. sturtevanti*, the synteny blocks were defined when all the genes were found in the same order and orientation in both species. The identified collinear blocks were then searched for the remaining genomes. Blocks with missing data, *i.e.* missing genes for one or more species were subsequently removed, and the remaining blocks were subjected to a size-based filtering with a threshold of ≥50kb. The selected blocks were aligned using the Mafft software (Katoh and Standley 2013).

### Pseudo-reference inference

Illumina reads were aligned on either the *D. willistoni* reference genome (Clark et al. 2007: 12) or the recently published *D. sturtevanti* assembly (Kim et al. 2021) using minimap2 (Li 2018). Minimap2 generated BAM files were assembled into a pseudo-reference for each species using the BEDTools software (Quinlan and Hall 2010). Each scaffold was then split into 100 kb-long windows, and phylogenetic analyses were conducted on each window separately using IqTree. A supertree was then inferred for each pseudo-reference genome approach using ASTRAL-III.

### Quantifying reticulation using the 2A2B test

Combinations of 28 species quartets were evaluated to test reticulation, bi-allelic sited shared by pairs were searched in every locus, whether using SCG, syntenic blocks or pseudo-reference 100 kb-long windows. Loci with at least 20 informative sites were retained. For each locus, the occurrences AABB, ABBA or BABA patterns were counted, and three pairwise χ^2^-based tests were conducted, denoted *D1*, *D2*, and *D3* as follows (D1 is reminiscent to Patterson’s D):

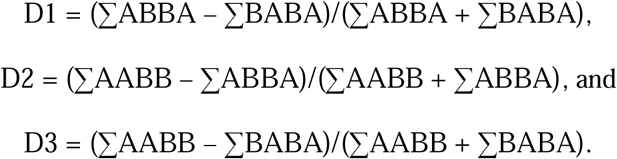

Each of these estimates is reminiscent to the classical Tajima’s (1993) Relative Rate Test (RTT), which tests the difference in branch lengths from two ingroup taxa to a third outgroup taxon, with lengths estimated as counts of different sites. However, unlike RRT, branch lengths in our analyses are counted in terms of shared differences, *i.e.* singletons are not considered. Similar to RRT, we applied a chi-squared analysis to test for deviation from parity for each parameter. According to the significance levels of the three estimates, a locus can be classified under each of the four categories presented in Figure 1. The test is run by the 2A2B perl script associated with this manuscript.

### Historical biogeography

To determine the sampling locations of the evaluated species, we conducted searches in TaxoDros (https://www.taxodros.uzh.ch/search/class.php). Additionally, we incorporated sampling sites that we ourselves had conducted. The accuracy of our species identification was confirmed through DNA barcoding. After inspection of the geographical points and manual correction, we identified the most northern, southern, western, and eastern points for each species. We used those points to reconstruct the ancestral geographical range by conducting a BayesTraits (Meade and Pagel 2022) analysis for each cartesian point separately. The analyses were carried out using the GEO model with the phylogenetic tree generated reconstructed with the Muller element A (XL chromosome arm), 1,000,000 of MCMC iterations and 25% burn-in. The divergence times were estimated using this tree under Bayesian inference. The calibration point used was the split between *D. willistoni* and *D. sturtevanti* at 17.5 myr, as estimated by Suvorov et al. (2022).

## Data availability statement

Genome sequences generated for this study are available on NCBI BioProject: PRJNA1078835.

## Acknowledgments

This paper is dedicated to Prof. Hermione E. M. C. Bicudo for her significant contributions to the study of the evolutionary genetics of species of the *Drosophila saltans* species group. We would like to thank David Ogereau for his assistance and insights in genome assembly for this study. We thank Fundação de Amparo à Pesquisa do Estado de São Paulo (FAPESP) for the financial support to L.M.R. (Number processes: 95/06165-1, 2014/14059-0 and 2016/ 11994-5) enabling us to collect and sequence many strains used here. We extend our thanks to the Conselho Nacional de Desenvolvimento Científico e Tecnológico (CNPq) for their support, specifically grant number 141545/2020-8, as well as the France Excellence Eiffel Scholarship Program for funding C.P.’s PhD scholarship. *Illumina* sequencing of *D. neocordata* was performed at the SNP&SEQ Technology Platform in Uppsala, Sweden, which is part of the Swedish National Genomics Infrastructure and Science for Life Laboratory. SNP&SEQ is supported by the Swedish Research Council and the Knut and Alice Wallenberg Foundation. This project was partly supported by a grant from The Swedish research council VR (2014-4353) to L.K. We want to thank the Nouragues research field station (managed by CNRS), which benefits from “Investissement d’Avenir” grants managed by Agence Nationale de la Recherche (AnaEE France ANR-11-INBS-0001; Labex CEBA ANR-10-LABX-25-01). The project was partly funded by the Austrian Science Fund FWF grant P28255-B22 to W.J.M. and by the Richard Lounsbery Foundation to A.Y.

## List of Supplementary Figures

**Figure S1 –** Phylogenetic relationships inferred using BEAST from BUSCO genes concatenated according to their Muller’s element.

**Figure S2 –** Phylogenetic relationships inferred using IqTree-2 from BUSCO genes concatenated according to their Muller’s element.

**Figure S3 –** Phylogenetic relationships inferred using ASTRAL-III from BUSCO genes concatenated according to their Muller’s element.

**Figure S4 –** Phylogenetic relationships inferred using ASTRAL-III from 100-kb long windows for pseudo-references produced by aligning short reads either to A) the ingroup *D. sturtevanti* or B) the outgroup *D. willistoni* reference assemblies.

## List of Supplementary Tables

**Table S1 –** A synopsis of phylogenetic relationships previously published in the *Drosophila saltans* species group. Species subgroups are represented by the first two letters in capital, whereas species names are represented by the first three letter in small.

**Table S2 –** Source and genome assembly statistics including busco scores for the genomes used in this study.

